# High-resolution image reconstruction with latent diffusion models from human brain activity

**DOI:** 10.1101/2022.11.18.517004

**Authors:** Yu Takagi, Shinji Nishimoto

## Abstract

Reconstructing visual experiences from human brain activity offers a unique way to understand how the brain represents the world, and to interpret the connection between computer vision models and our visual system. While deep generative models have recently been employed for this task, reconstructing realistic images with high semantic fidelity is still a challenging problem. Here, we propose a new method based on a diffusion model (DM) to reconstruct images from human brain activity obtained via functional magnetic resonance imaging (fMRI). More specifically, we rely on a latent diffusion model (LDM) termed Stable Diffusion. This model reduces the computational cost of DMs, while preserving their high generative performance. We also characterize the inner mechanisms of the LDM by studying how its different components (such as the latent vector of image Z, conditioning inputs C, and different elements of the denoising U-Net) relate to distinct brain functions. We show that our proposed method can reconstruct high-resolution images with high fidelity in straight-forward fashion, without the need for any additional training and fine-tuning of complex deep-learning models. We also provide a quantitative interpretation of different LDM components from a neuroscientific perspective. Overall, our study proposes a promising method for reconstructing images from human brain activity, and provides a new framework for understanding DMs. Please check out our webpage at https://sites.google.com/view/stablediffusion-with-brain/.

## 1. Introduction

A fundamental goal of computer vision is to construct artificial systems that see and recognize the world as human visual systems do. Recent developments in the measurement of population brain activity, combined with advances in the implementation and design of deep neural network models, have allowed direct comparisons between latent representations in biological brains and architectural characteristics of artificial networks, providing important insights into how these systems operate [3, 8–10, 13, 18, 19, 21, 42, 43, 54, 55]. These efforts have included the reconstruction of visual experiences (perception or imagery) from brain activity, and the examination of potential correspondences between the computational processes associated with biological and artificial systems [2, 5, 7, 24, 25, 27, 36, 44–46].

Reconstructing visual images from brain activity, such as that measured by functional Magnetic Resonance Imaging (fMRI), is an intriguing but challenging problem, because the underlying representations in the brain are largely unknown, and the sample size typically associated with brain data is relatively small [17, 26, 30, 32]. In recent years, researchers have started addressing this task using deep-learning models and algorithms, including generative adversarial networks (GANs) and self-supervised learning [2, 5, 7, 24, 25, 27, 36, 44–46]. Additionally, more recent studies have increased semantic fidelity by explicitly using the semantic content of images as auxiliary inputs for reconstruction [5, 25]. However, these studies require training new generative models with fMRI data from scratch, or fine-tuning toward the specific stimuli used in the fMRI experiment. These efforts have shown impressive but limited success in pixel-wise and semantic fidelity, partly because the number of samples in neuroscience is small, and partly because learning complex generative models poses numerous challenges.

Diffusion models (DMs) [11,47,48,53] are deep generative models that have been gaining attention in recent years. DMs have achieved state-of-the-art performance in several tasks involving conditional image generation [4,39,49], image super resolution [40], image colorization [38], and other related tasks [6, 16, 33, 41]. In addition, recently proposed latent diffusion models (LDMs) [37] have further reduced computational costs by utilizing the latent space generated by their autoencoding component, enabling more efficient computations in the training and inference phases. Another advantage of LDMs is their ability to generate high-resolution images with high semantic fidelity. However, because LDMs have been introduced only recently, we still lack a satisfactory understanding of their internal mechanisms. Specifically, we still need to discover how they represent latent signals within each layer of DMs, how the latent representation changes throughout the denoising process, and how adding noise affects conditional image generation.

Here, we attempt to tackle the above challenges by reconstructing visual images from fMRI signals using an LDM named Stable Diffusion. This architecture is trained on a large dataset and carries high text-to-image generative performance. We show that our simple framework can reconstruct high-resolution images with high semantic fidelity without any training or fine-tuning of complex deeplearning models. We also provide biological interpretations of each component of the LDM, including forward/reverse diffusion processes, U-Net, and latent representations with different noise levels.

Our contributions are as follows: (i) We demonstrate that our simple framework can reconstruct high-resolution (512 × 512) images from brain activity with high semantic fidelity, without the need for training or fine-tuning of complex deep generative models (Figure 1); (ii) We quantitatively interpret each component of an LDM from a neuroscience perspective, by mapping specific components to distinct brain regions; (iii) We present an objective interpretation of how the text-to-image conversion process implemented by an LDM incorporates the semantic information expressed by the conditional text, while at the same time maintaining the appearance of the original image.

**Figure 1.**
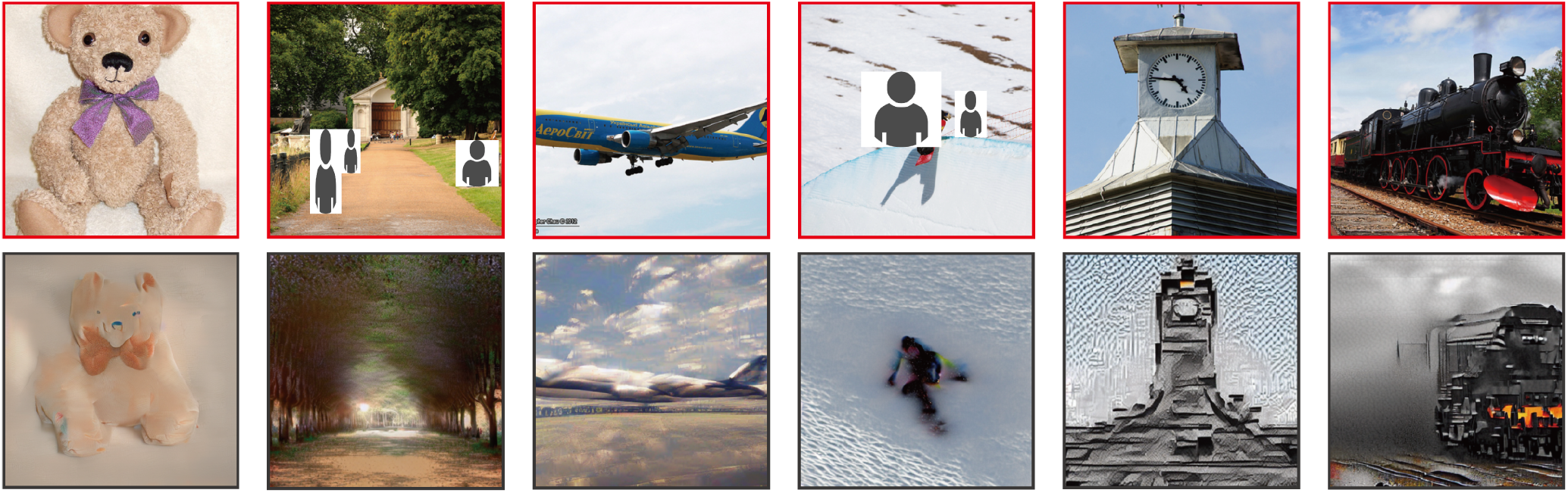
Presented images (red box, top row) and images reconstructed from fMRI signals (gray box, bottom row) for one subject (subj01).

## 2. Related Work

### 2.1. Reconstructing visual image from fMRI

Decoding visual experiences from fMRI activity has been studied in various modalities. Examples include explicitly presented visual stimuli [17, 26, 30, 32], semantic content of the presented stimuli [15, 31, 52], imagined content [13, 29], perceived emotions [12, 20, 51], and many other related applications [14, 28]. In general, these decoding tasks are made difficult by the low signal-to-noise ratio and the relatively small sample size associated with fMRI data.

While early attempts have used handcrafted features to reconstruct visual images from fMRI [17,26,30,32], recent studies have begun to use deep generative models trained on a large number of naturalistic images [2, 5, 7, 24, 25, 27, 36, 44–46]. Additionally, a few studies have used semantic information associated with the images, including categorical or text information, to increase the semantic fidelity of the reconstructed images [5, 25]. To produce high-resolution reconstructions, these studies require training and possibly fine-tuning of generative models, such as GANs, with the same dataset used in the fMRI experiments. These requirements impose serious limitations, because training complex generative models is in general challenging, and the number of samples in neuroscience is relatively small. Thus, even modern implementations struggle to produce images, at most 256 × 256 resolution, with high semantic fidelity unless they are augmented with numerous tools and techniques. DMs and LDMs are recent algorithms for image generation that could potentially address these limitations, thanks to their ability to generate diverse high-resolution images with high semantic fidelity of text-conditioning, and high computational efficiency. However, to the best of our knowledge, no prior studies have used DMs for visual reconstruction.

### 2.2. Encoding Models

To understand deep-learning models from a biological perspective, neuroscientists have employed encoding models: a predictive model of brain activity is built out of features extracted from different components of the deeplearning models, followed by examination of the potential link between model representations and corresponding brain processes [3, 8–10, 13, 18, 19, 21, 42, 43, 54, 55]. Because brains and deep-learning models share similar goals (e.g., recognition of the world) and thus could implement similar functions, the ability to establish connections between these two structures provides us with biological interpretations of the architecture underlying deep-learning models, otherwise viewed as black boxes. For example, the activation patterns observed within early and late layers of a CNN correspond to the neural activity patterns measured from early and late layers of visual cortex, suggesting the existence of a hierarchical correspondence between latent representations of a CNN and those present in the brain [9, 10, 13, 19, 54, 55]. This approach has been applied primarily to vision science, but it has recently been extended to other sensory modalities and higher functions [3, 8, 18, 21, 42, 43].

Compared with biologically inspired architectures such as CNNs, the correspondence between DMs and the brain is less obvious. By examining the relationship between each component and process of DMs and corresponding brain activities, we were able to obtain biological interpretations of DMs, for example in terms of how latent vectors, denoising processes, conditioning operations, and U-net components may correspond to our visual streams. To our knowledge, no prior study has investigated the relationship between DMs and the brain.

Together, our overarching goal is to use DMs for high resolution visual reconstruction and to use brain encoding framework to better understand the underlying mechanisms of DMs and its correspondence to the brain.

## 3. Methods

Figure 2 presents an overview of our methods.

**Figure 2.**
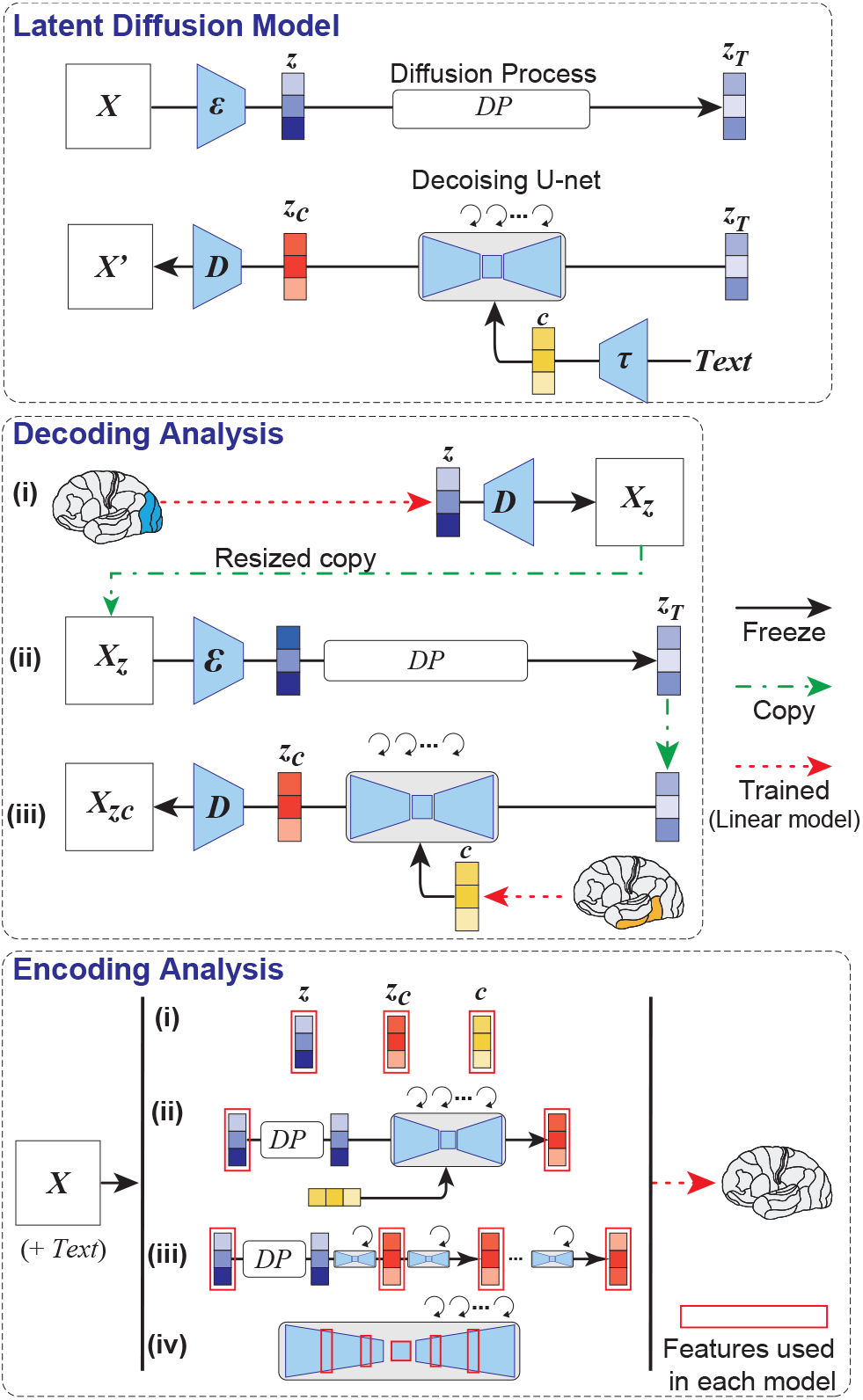
Overview of our methods. (Top) Schematic of LDM used in this study. *ε* denotes an image encoder, *D* is a image decoder, and *τ* is a text encoder (CLIP). (Middle) Schematic of decoding analysis. We decoded latent representations of the presented image (z) and associated text c from fMRI signals within early (blue) and higher (yellow) visual cortices, respectively. These latent representations were used as input to produce a reconstructed image X_zc_. (Bottom) Schematic of encoding analysis. We built encoding models to predict fMRI signals from different components of LDM, including z, c, and z_c_.

### 3.1. Dataset

We used the Natural Scenes Dataset (NSD) for this project [1]. Please visit the NSD website for more details ^1^. Briefly, NSD provides data acquired from a 7-Tesla fMRI scanner over 30–40 sessions during which each subject viewed three repetitions of 10,000 images. We analyzed data for four of the eight subjects who completed all imaging sessions (subj01, subj02, subj05, and subj07). The images used in the NSD experiments were retrieved from MS COCO and cropped to 425 × 425 (if needed). We used 27,750 trials from NSD for each subject (2,250 trials out of the total 30,000 trials were not publicly released by NSD). For a subset of those trials (N=2,770 trials), 982 images were viewed by all four subjects. Those trials were used as the test dataset, while the remaining trials (N=24,980) were used as the training dataset.

For functional data, we used the preprocessed scans (resolution of 1.8 mm) provided by NSD. See Appendix A for details of the preprocessing protocol. We used single-trial beta weights estimated from generalized linear models and region of interests (ROIs) for early and higher (ventral) visual regions provided by NSD. For the test dataset, we used the average of the three trials associated with each image. For the training dataset, we used the three separate trials without averaging.

### 3.2. Latent Diffusion Models

DMs are probabilistic generative models that restore a sampled variable from Gaussian noise to a sample of the learned data distribution via iterative denoising. Given training data, the diffusion process destroys the structure of the data by gradually adding Gaussian noise. The sample at each time point is defined as 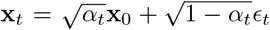 where x_*t*_ is a noisy version of input x_0_, *t* ∈ {1,…, *T*}, *α* is a hyperparameter, and *ε* is the Gaussian. The inverse diffusion process is modeled by applying a neural network *f_θ_*(x_*t*_, *t*) to the samples at each step to recover the original input. The learning objective is *f_θ_*(x, *t*) ≈ *ε_t_* [11, 47]. U-Net is commonly used for neural networks *f_θ_*.

This method can be generalized to learning conditional distributions by inserting auxiliary input c into the neural network. If we set the latent representation of the text sequence to c, it can implement text-to-image models. Recent studies have shown that, by using large language and image models, DMs can create realistic, high-resolution images from text inputs. Furthermore, when we start from source image with input texts, we can generate new text conditional images by editing the image. In this image-to-image translation, the degree of degradation from the original image is controlled by a parameter that can be adjusted to preserve either the semantic content or the appearance of the original image.

DMs that operate in pixel space are computationally expensive. LDMs overcome this limitation by compressing the input using an autoencoder (Figure 2, top). Specifically, the autoencoder is first trained with image data, and the diffusion model is trained to generate its latent representation z using a U-Net architecture. In doing so, it refers to conditional inputs via cross-attention. This allows for lightweight inference compared with pixel-based DMs, and for very high-quality text-to-image and image-to-image implementations.

In this study, we used an LDM called Stable Diffusion, which was built on LDMs and trained on a very large dataset. The model can generate and modify images based on text input. Text input is projected to a fixed latent representation by a pretrained text encoder (CLIP) [34]. We used version 1.4 of the model. See Appendix A for details on the training protocol.

We define z as the latent representation of the original image compressed by the autoencoder, c as the latent representation of texts (average of five text annotations associated to each MS COCO image), and z_c_ as the generated latent representation of z modified by the model with c. We used these representations in the decoding/encoding models described below.

### 3.3. Decoding: reconstructing images from fMRI

We performed visual reconstruction from fMRI signals using LDM in three simple steps as follows (Figure 2, middle). The only training required in our method is to construct linear models that map fMRI signals to each LDM component, and no training or fine-tuning of deep-learning models is needed. We used the default parameters of image-to-image and text-to-image codes provided by the authors of LDM ^2^, including the parameters used for the DDIM sampler. See Appendix A for details.

i. First, we predicted a latent representation z of the presented image X from fMRI signals within early visual cortex. z was then processed by an decoder of autoencoder to produce a coarse decoded image X_z_ with a size of 320 × 320, and then resized it to 512 × 512.
ii. X_z_ was then processed by encoder of autoencoder, then added noise through the diffusion process.
iii. We decoded latent text representations c from fMRI signals within higher (ventral) visual cortex. Noise-added latent representations z_T_ of the coarse image and decoded c were used as input to the denoising U-Net to produce z_c_. Finally, z_c_ was used as input to the decoding module of the autoencoder to produce a final reconstructed image X_zc_ with a size of 512 × 512.

To construct models from fMRI to the components of LDM, we used L2-regularized linear regression, and all models were built on a per subject basis. Weights were estimated from training data, and regularization parameters were explored during the training using 5-fold crossvalidation. We resized original images from 425 × 425 to 320 × 320 but confirmed that resizing them to a larger size (448 × 448) does not affect the quality of reconstruction.

As control analyses, we also generated images using only z or c. To generate these control images, we simply omitted c or z from step (iii) above, respectively.

The accuracy of image reconstruction was evaluated objectively (perceptual similarity metrics, psms) and subjectively (human raters, N=6) by assessing whether the original test images (N=982 images) could be identified from the generated images. As a similarity metrics of PSMs, we used early/middle/late layers of CLIP and CNN (AlexNet) [22]. Briefly, we conducted two-way identification experiments: examined whether the image reconstructed from fMRI was more similar to the corresponding original image than randomly picked reconstructed image. See Appendix B for details and additional results.

### 3.4. Encoding: Whole-brain Voxel-wise Modeling

Next, we tried to interpret the internal operations of LDMs by mapping them to brain activity. For this purpose, we constructed whole-brain voxel-wise encoding models for the following four settings (see Figure 2 bottom and Appendix A for implementation details):

i. We first built linear models to predict voxel activity from the following three latent representations of the LDM independently: z, c, and z_c_.
ii. Although z_c_ and z produce different images, they result in similar prediction maps on the cortex (see 4.2.1). Therefore, we incorporated them into a single model, and further examined how they differ by mapping the unique variance explained by each feature onto cortex [23]. To control the balance between the appearance of the original image and the semantic fidelity of the conditional text, we varied the level of noise added to z. This analysis enabled quantitative interpretation of the image-to-image process.
iii. While LDMs are characterized as an iterative denoising process, the internal dynamics of the denoising process are poorly understood. To gain some insight into this process, we examined how z_c_ changes through the denoising process. To do so, we extracted z_c_ from the early, middle, and late steps of the denoising. We then constructed combined models with z as in the above analysis (ii), and mapped their unique variance onto cortex.
iv. Finally, to inspect the last black box associated with LDMs, we extracted features from different layers of U-Net. For different steps of the denoising, encoding models were constructed independently with different U-Net layers: two from the first stage, one from the bottleneck stage, and two from the second stage. We then identified the layer with highest accuracy for each voxel and for each step.

Model weights were estimated from training data using L2-regularized linear regression, and subsequently applied to test data (see Appendix A for details). For evaluation, we used Pearson’s correlation coefficients between predicted and measured fMRI signals. We computed statistical significance (one-sided) by comparing the estimated correlations to the null distribution of correlations between two independent Gaussian random vectors of the same length (N=982). The statistical threshold was set at *P* < 0.05 and corrected for multiple comparisons using the FDR procedure. We show results from a single random seed, but we verified that different random seed produced nearly identical results (see Appendix C). We reduced all feature dimensions to 6,400 by applying principal component analysis, by estimating components within training data.

## 4. Results

### 4.1. Decoding

Figure 3 shows the results of visual reconstruction for one subject (subj01). We generated five images for each test image and selected the generated images with highest PSMs. On the one hand, images reconstructed using only z were visually consistent with the original images, but failed to capture their semantic content. On the other hand, images reconstructed using only c generated images with high semantic fidelity but were visually inconsistent. Finally, images reconstructed using z_c_ could generate high-resolution images with high semantic fidelity (see Appendix B for more examples).

**Figure 3.**
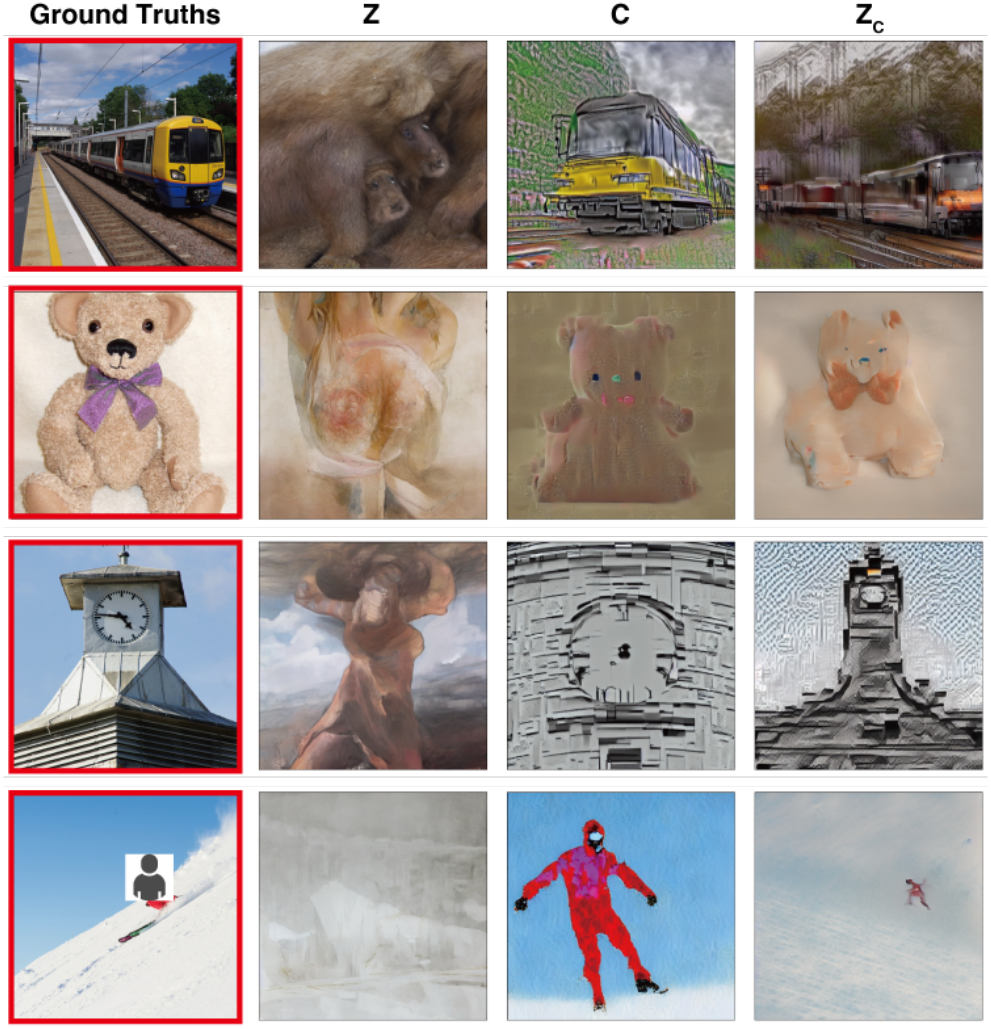
Presented (red box) and reconstructed images for a single subject (subj01) using z, c, and z_c_.

Figure 4 shows reconstructed images from all subjects for the same image (all images were generated using z_c_. Other examples are available in the Appendix B). Overall, reconstruction quality was stable and accurate across subjects.

**Figure 4.**
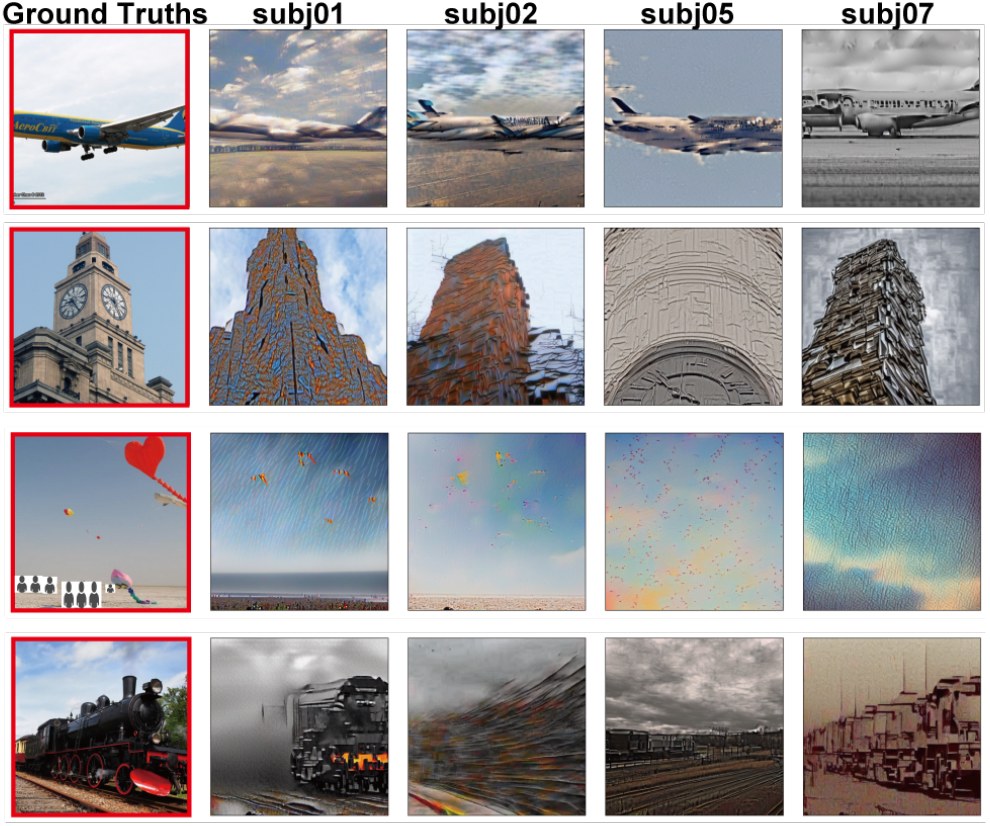
Example results for all four subjects.

We note that, the lack of agreement regarding specific details of the reconstructed images may differences in perceived experience across subjects, rather than failures of reconstruction. Alternatively it may simply reflect differences in data quality among subjects. Indeed, subjects with high (subj01) and low (subj07) decoding accuracy from fMRI were subjects with high and low data quality metrics, respectively (see Appendix B).

Figure 5 plots results for the quantitative evaluation. In the objective evaluation, images reconstructed using z_c_ are generally associated with higher accuracy values across different metrics than images reconstructed using only z or c. When only z was used, accuracy values were particularly high for PSMs derived from early layers of CLIP and CNN. On the other hand, when only c was used, accuracy values were higher for PSMs derived from late layers. In the subjective evaluation, accuracy values of images obtained from c are higher than those obtained from z, while z_c_ resulted in the highest accuracy compared with the other two methods (*P* < 0.01 for all comparisons, two-sided signed-rank test, FWE corrected). Together, these results suggest that our method captures not only low-level visual appearance, but also high-level semantic content of the original stimuli.

**Figure 5.**
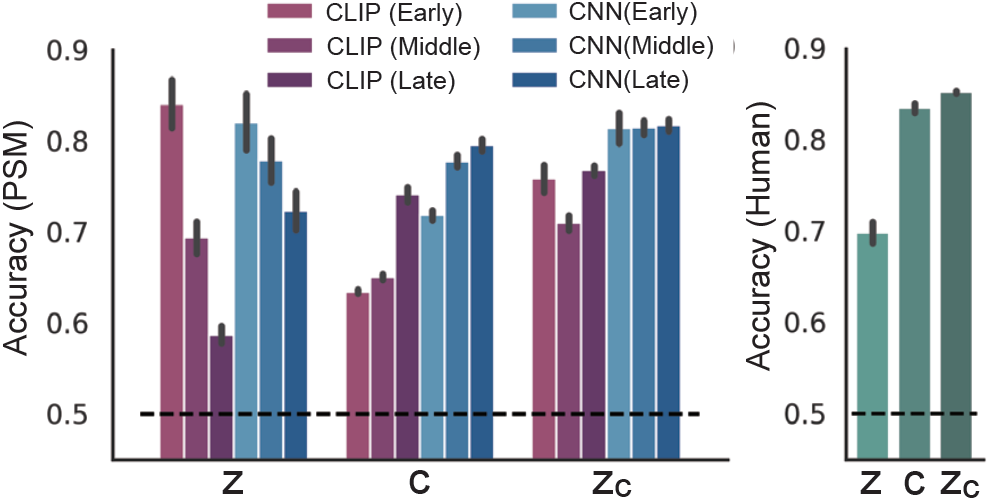
Identification accuracy calculated using objective (left) and subjective (right) criteria (pooled across four subjects; chance level corresponds to 50%). Error bars indicate standard error of the mean.

It is difficult to compare our results with those reported by most previous studies, because they used different datasets. The datasets used in previous studies contain far fewer images, much less image complexity (typically individual objects positioned in the center of the image), and lack full-text annotations of the kind available from NSD. Only one study to date [25] used NSD for visual reconstruction, and they reported accuracy values of 78 ±4.5% for one subject (subj01) using PSM based on Inception V3. It is difficult to draw a direct comparison with this study, because it differed from ours in several respects (for example, it used different training and test sample sizes, and different image resolutions). Notwithstanding these differences, their reported values fall within a similar range to ours for the same subject (77% using CLIP, 83% using AlexNet, and 76% using Inception V3). However, this prior study relied on extensive model training and feature engineering with many more hyper-parameters than those adopted in our study, including the necessity to train complex generative models, fine-tuning toward MS COCO, data augmentation, and arbitrary thresholding of features. We did not use any of the above techniques — rather, our simple pipeline only requires the construction of two linear regression models from fMRI activity to latent representations of LDM.

Furthermore, we observed a reduction in semantic fidelity when we used categorical information associated with the images, rather than full-text annotations for c. We also found an increase in semantic fidelity when we used semantic maps instead of original images for z, though visual similarity was decreased in this case (see Appendix B).

### 4.2. Encoding Model

#### 4.2.1 Comparison among Latent Representations

Figure 6 shows prediction accuracy of the encoding models for three types of latent representations associated with the LDM: z, a latent representation of the original image; c, a latent representation of image text annotation; and z_c_, a noise-added latent representation of z after reverse diffusion process with cross-attention to c.

**Figure 6.**
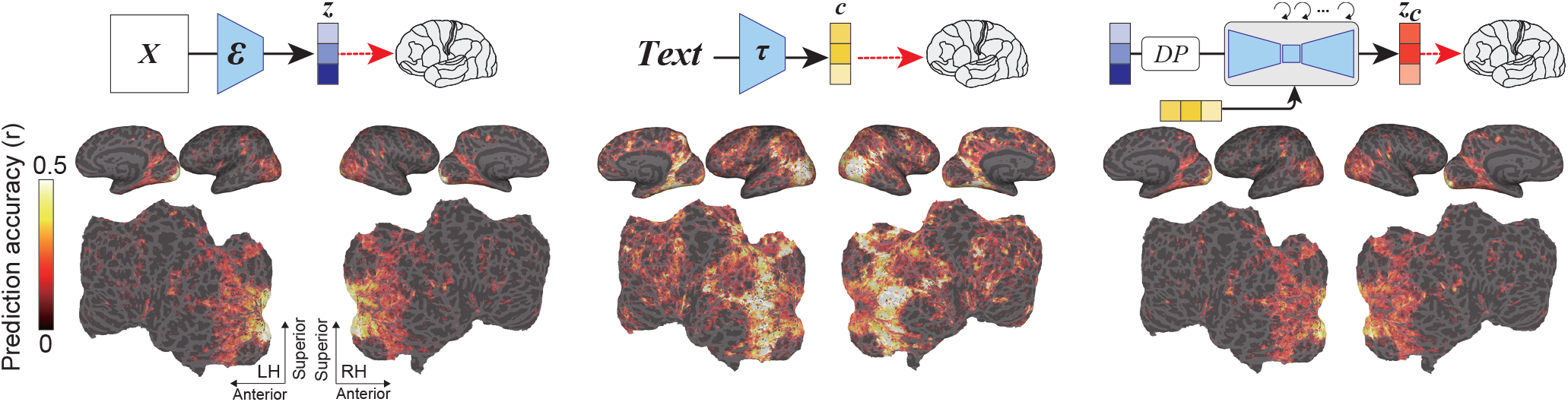
Prediction performance (measured using Pearson’s correlation coefficients) for the voxel-wise encoding model applied to held-out test images in a single subject (subj01), projected onto the inflated (top, lateral and medial views) and flattened cortical surface (bottom, occipital areas are at the center), for both left and right hemispheres. Brain regions with significant accuracy are colored (all colored voxels *P* < 0.05, FDR corrected).

Although all three components produced high prediction performance at the back of the brain, visual cortex, they showed stark contrast. Specifically, z produced high prediction performance in the posterior part of visual cortex, namely early visual cortex. It also showed significant prediction values in the anterior part of visual cortex, namely higher visual cortex, but smaller values in other regions. On the other hand, c produced the highest prediction performance in higher visual cortex. The model also showed high prediction performance across a wide span of cortex. z_c_ carries a representation that is very similar to z, showing high prediction performance for early visual cortex. Although this is somewhat predictable given their intrinsic similarity, it is nevertheless intriguing because these representations correspond to visually different generated images. We also observed that using z_c_ with a reduced noise level injected into z produces a more similar prediction map to the prediction map obtained from z, as expected (see Appendix C). This similarity prompted us to conduct an additional analysis to compare the unique variance explained by these two models, detailed in the following section. See Appendix C for results of all subjects.

#### 4.2.2 Comparison across different noise levels

While the previous results showed that prediction accuracy maps for z and z_c_ present similar profiles, they do not tell us how much unique variance is explained by each feature as a function of different noise levels. To enhance our understanding of the above issues, we next constructed encoding models that simultaneously incorporated both z and z_c_ into a single model, and studied the unique contribution of each feature. We also varied the level of noise added to z for generating z_c_.

Figure 7 shows that, when a small amount of noise was added, z predicted voxel activity better than z_c_ across cortex. Interestingly, when we increased the level of noise, z_c_ predicted voxel activity within higher visual cortex better than z, indicating that the semantic content of the image was gradually emphasized.

**Figure 7.**
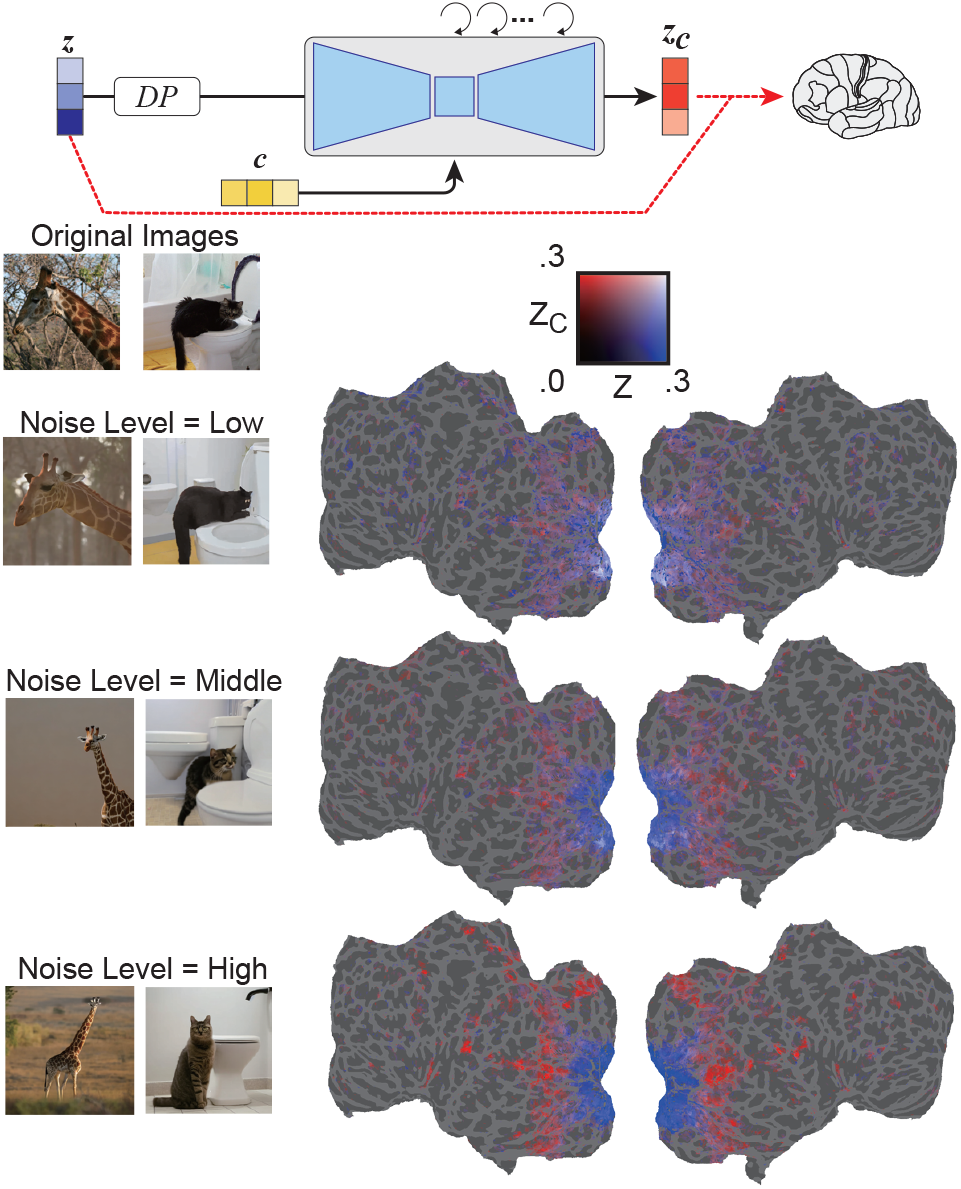
Unique variance accounted for by z_c_ compared with z in one subject (subj01), obtained by splitting accuracy values from the combined model. While fixing z, we used z_c_ with varying amounts of noise-level added to the latent representation of stimuli from low-level (top) to high-level (bottom). All colored voxels *P* < 0.05, FDR corrected.

This result is intriguing because, without analyses like this, we can only observe randomly generated images, and we cannot examine how the text-conditioned image-to-image process is able to balance between semantic content and original visual appearance.

#### 4.2.3 Comparison across different diffusion stages

We next asked how the noise-added latent representation changes over the iterative denoising process.

Figure 8 shows that, during the early stages of the denoising process, z signals dominated prediction of fMRI signals. During the middle step of the denoising process, z_c_ predicted activity within higher visual cortex much better than z, indicating that the bulk of the semantic content emerges at this stage. These results show how LDM refines and generates images from noise.

**Figure 8.**
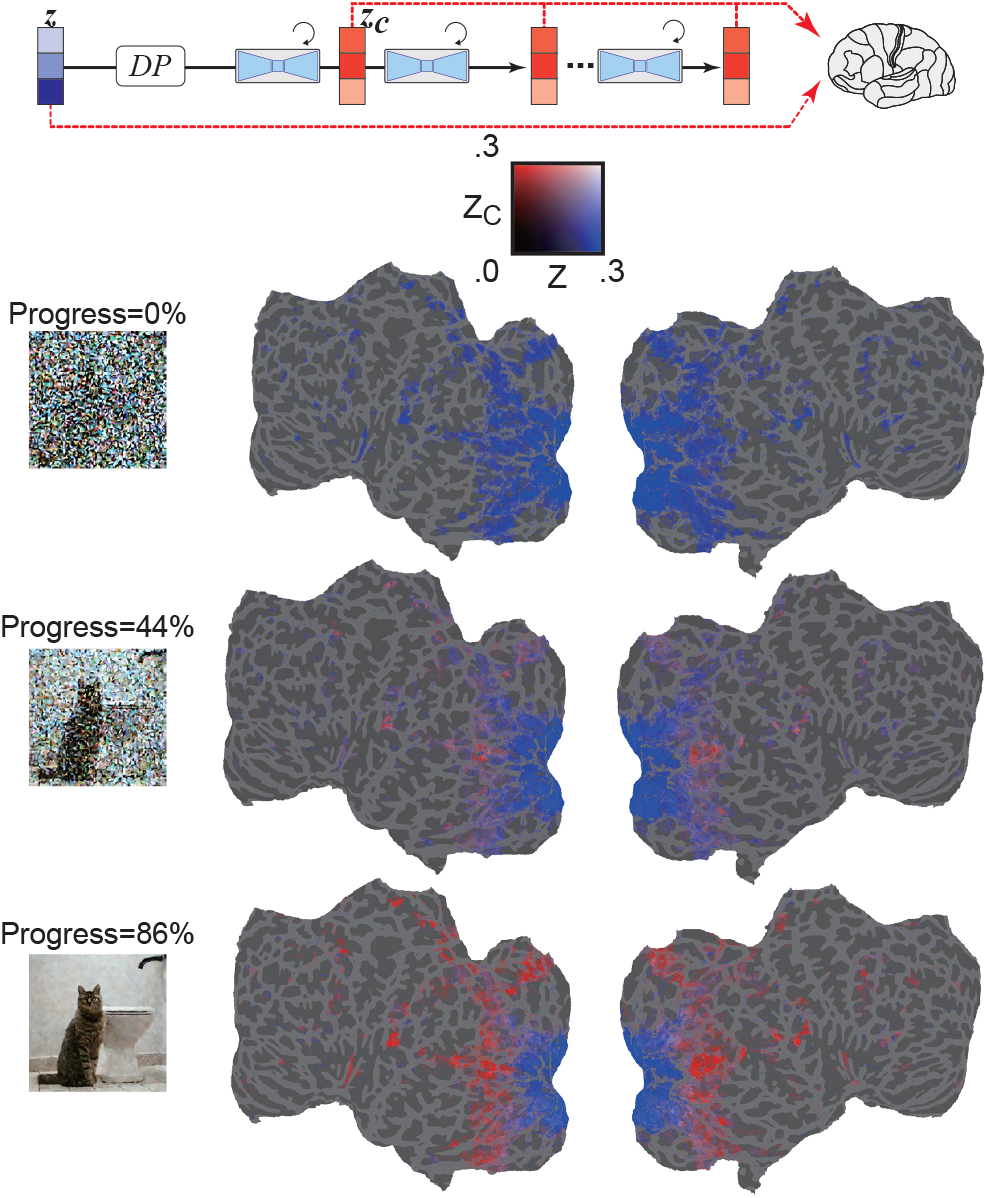
Unique variance accounted for by z_c_ compared with z in one subject (subj01), obtained by splitting accuracy values from the combined model. While fixing z, we used z_c_ with different denoising stages from early (top) to late (bottom) steps. All colored voxels *P* < 0.05, FDR corrected.

#### 4.2.4 Comparison across different U-Net Layers

Finally, we asked what information is being processed at each layer of U-Net.

Figure 9 shows the results of encoding models for different steps of the denoising process (early, middle, late), and for the different layers of U-Net. During the early phase of the denoising process, the bottleneck layer of U-Net (colored orange) produces the highest prediction performance across cortex. However, as denoising progresses, the early layer of U-Net (colored blue) predicts activity within early visual cortex, and the bottleneck layer shifts toward superior predictive power for higher visual cortex.

**Figure 9.**
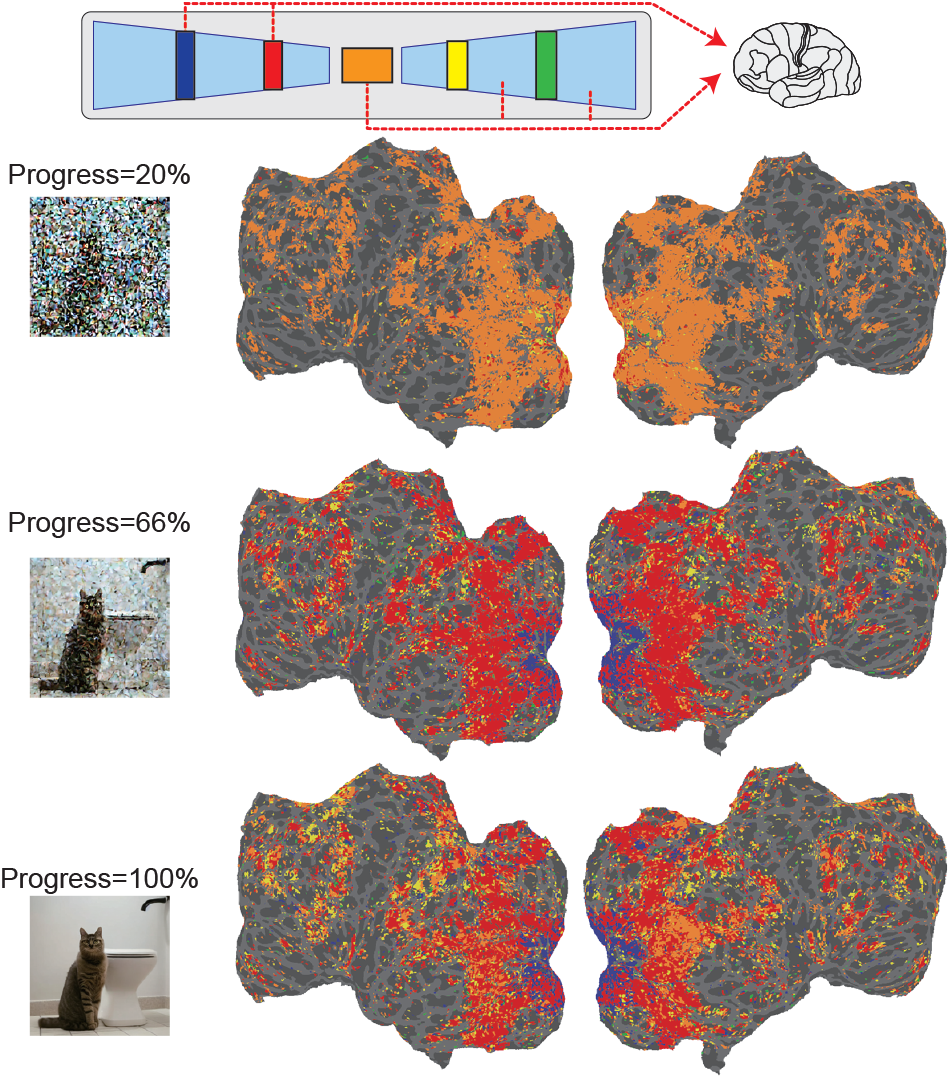
Selective engagement of different U-Net layers for different voxels across the brain. Colors represent the most predictive U-Net layer for early (top) to late (bottom) denoising steps. All colored voxels *P* < 0.05, FDR corrected.

These results suggest that, at the beginning of the reverse diffusion process, image information is compressed within the bottleneck layer. As denoising progresses, a functional dissociation among U-Net layers emerges within visual cortex: i.e., the first layer tends to represent fine-scale details in early visual areas, while the bottleneck layer corresponds to higher-order information in more ventral, semantic areas.

## 5. Conclusions

We propose a novel visual reconstruction method using LDMs. We show that our method can reconstruct high-resolution images with high semantic fidelity from human brain activity. Unlike previous studies of image reconstruction, our method does not require training or fine-tuning of complex deep-learning models: it only requires simple linear mappings from fMRI to latent representations within LDMs.

We also provide a quantitative interpretation for the internal components of the LDM by building encoding models. For example, we demonstrate the emergence of semantic content throughout the inverse diffusion process, we perform layer-wise characterization of U-Net, and we provide a quantitative interpretation of image-to-image transformations with different noise levels. Although DMs are developing rapidly, their internal processes remain poorly understood. This study is the first to provide a quantitative interpretation from a biological perspective.

## Acknowledgements

We would like to thank Computer Vision and Learning research group at Ludwig Maximilian University of Munich for providing the codes and models for Stable Diffusion, Stability AI for supporting the training of Stable Diffusion, and NSD for providing the neuroimaging dataset. YT was supported by JSPS KAKENHI (19H05725). SN was supported by MEXT/JSPS KAKENHI JP18H05522 as well as JST CREST JPMJCR18A5 and ERATO JPMJER1801.

## Footnotes

1 http://naturalscenesdataset.org/

2 https://github.com/CompVis/stable-diffusion/blob/main/scripts/

## Notes

https://sites.google.com/view/stablediffusion-with-brain/

### Competing Interest Statement

The authors have declared no competing interest.

### Summary of Updates

Modified Acknowledgement. Other than that, nothing has been changed.

