## Appendix for "High-resolution image reconstruction with latent diffusion models from human brain activity"

### A. Additional Method Details

#### A.1. Latent Diffusion Models

We used the Stable Diffusion model released by Computer Vision and Learning research group at Ludwig Maximilian University of Munich. Stable Diffusion is a generative model that generates text-conditional images (text-to-image). It also performs other tasks, such as image translation (image-to-image). This model is trained on $512 \times 512$ images from a subset of the LAION-5B database. It uses a frozen CLIP ViT-L/14 text encoder to condition the model on text prompts. The model is lightweight compared with previous diffusion models and runs on a GPU with at least 10 GB VRAM. We adopted version 1.4 of the model provided by Hugging Face.

We used the default parameters of image-to-image code provided by the authors of the latent diffusion model (LDM) [3]. Specifically, we used the DDIM sampler with 50 sampling steps. We set the unconditional guidance scale parameter to 5.0. Additionally, we set the strength parameter for noising/denoising to 0.8, except for the encoding models with different noise levels.

#### A.2. Dataset

We used the preprocessed scans (with resolution of 1.8 mm) provided by Natural Scenes Dataset (NSD). Preprocessing of the functional data included temporal resampling, which corrected for differences in slice time acquisition, and spatial interpolation, which corrected for head motion and spatial distortion. NSD provides three types of single-trial beta weight estimated from generalized linear models. In this study, we used the version named *betasfithrfGLMdenoiseRR*. The beta weights were z-scored across runs separately for each voxel in each subject. NSD also provides several regions of interests (ROIs) that were identified using separate functional localization experiments in each subject. We used ROIs for early and higher (ventral) visual regions included in the *streams* atlas. For the test dataset, we used the average of the three trials associated with each image. For the training dataset, we used the three separate trials without averaging.

#### A.3. Decoding and Encoding Analysis

Figure A.1 is a schematic of the training pipeline for decoding and encoding analyses. For both analyses, we estimated weights of linear models using the training dataset. We then applied these weights to the test dataset, resulting in the prediction for features (decoding) or voxels (encoding). We used the predicted features for visual reconstruction analysis in the decoding analysis and predicted voxel activities for calculating Pearson's correlation coefficients between predicted and true voxel activities in the encoding analysis.

In the encoding analyses, we varied noise levels by changing the strength parameter for noising/denoising. Specifically, for Figure 7, we set the strength parameter to 0.6 (low), 0.8 (middle) and 0.9 (high). For Figures 6, 8 and 9, we set the parameter to 0.9.

In Figure 8 and 9, we varied the denoising steps to obtain $\mathbf{z_c}$ or U-net features. In the case of Figure 8, we extracted $\mathbf{z_c}$ from the outputs of U-net after 0, 20, and 40 denoising steps. In the case of Figure 9, we extracted features from each U-net layer after 10, 30, and all denoising steps. We used outputs (before downsample/upsample) of U-net from the second and third blocks of the first stage (input blocks), output of the bottleneck stage (middle block), and second and third blocks of the second stage (output blocks).

### B. Additional Results of Reconstruction

#### B.1. Decoding Accuracy of Latent Representations

We present numerical results for the decoding performance of $\mathbf{z}$ and $\mathbf{c}$ from fMRI signals in Table B.1. In this table, we also give two quality metrics provided by NSD, namely the temporal signal-to-noise ratio (tSNR) and $R^2$ values of the simple ON–OFF generalized linear model (GLM) fitted to the NSD data. A subject with high quality metrics (subj01) tended to have higher decoding accuracy.

#### B.2. Results of Identification

In the objective evaluation, for each original image, we generated five images with different stochastic noise. We then calculated the average similarity between the generated images and the corresponding original image. The identification

[3] https://github.com/CompVis/stable-diffusion/blob/main/scripts/

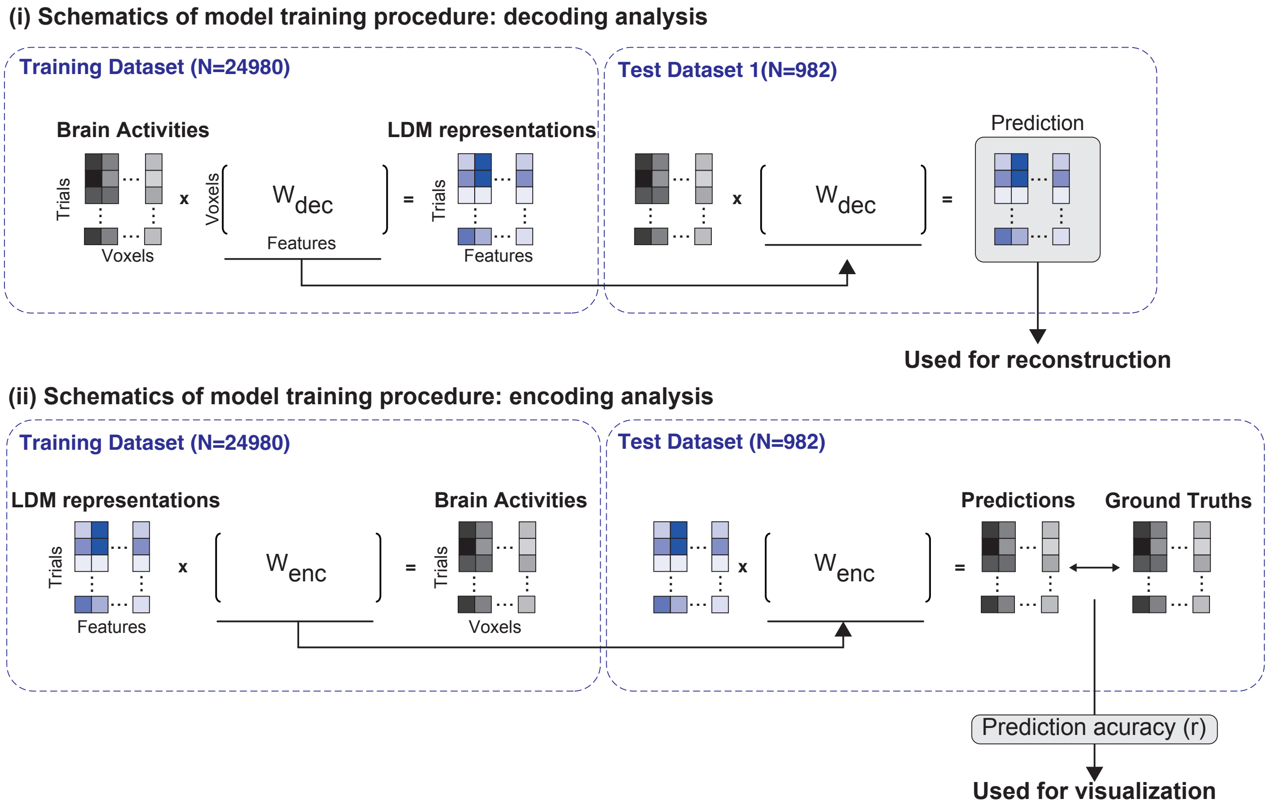


Figure A.1. Schematic of decoding and encoding analyses

Table B.1. Decoding accuracy (Pearson's correlation coefficients) of latent representations and data quality metrics. Mean±s.e.m. across all features are shown for **z** and **c**. Mean±s.t.d. across all sessions are shown for tSNR and the simple GLM.

| | | **subj01** | **subj02** | **subj05** | **subj07** |
|---|---|---|---|---|---|
| **Decoding accuracy** | **z** | 0.239 ± 0.137 | 0.213 ± 0.132 | 0.177 ± 0.134 | 0.145 ± 0.136 |
| | **c** | 0.304 ± 0.108 | 0.295 ± 0.107 | 0.341 ± 0.109 | 0.296 ± 0.118 |
| **Quality metrics (from NSD)** | **tSNR** | 42.09 ± 2.89 | 37.89 ± 3.58 | 35.70 ± 4.93 | 37.00 ± 6.74 |
| | **Simple GLM(ON-OFF) $R^2$** | 5.92 ± 2.12 | 4.08 ± 1.56 | 5.06 ± 2.03 | 3.57 ± 1.78 |

accuracy for each set of the generated images was defined as the winning rate of the true generated images compared with the other set of generated images. As similarity metrics, we used perceptual similarity metrics (PSMs) [35] computed via Pearson's coefficients of correlation between two images with reference to high-level image features. Here, we used early/middle/late layers of CLIP (ViT-L/14, Hugging Face implementation) and CNN (AlexNet, PyTorch implementation) [22]. Specifically, for CLIP, we used outputs from the 7th, 12th, and final layers of the vision encoder. For AlexNet, we used outputs of the rectified linear function before max-pooling (if any) from the second, fifth, and seventh layers. In Figure B.2, we also show the results when we used the feature before the last FC layer of Inception V3 [50] (PyTorch implementation).

For the subjective evaluation, we presented the original image, the corresponding generated image, and the randomly chosen non-corresponding generated image to six raters. The raters were asked to choose the generated image that appeared to be most similar to the original image. The answer was considered correct if the corresponding generated image was chosen. We presented three types of generated image: images generated from $\mathbf{z}$, $\mathbf{c}$, and $\mathbf{z_c}$. For each type of image, 800 images were shown to each rater (200 images $\times$ 4 subjects), and 2400 images were thus shown in total.

In Figure B.2, we present identification results with other types of features ($\mathbf{z}_{raw}$ and $\mathbf{z}_{cat}$) and PSM metrics (Inception V3). $\mathbf{z}_{raw}$ relates to the identification when we simply used the decoded image from the predicted $\mathbf{z}$ without forward/reverse diffusion. Although the accuracy is high for the low-level visual features, the high-level semantics of the original images were not captured. $\mathbf{z}_{cat}$ relates to the identification when we used only category information presented in the image, rather than full-text annotations for $\mathbf{c}$. We observed a reduction in semantic fidelity when we used only the categorical information.

In Figure B.3, we present identification results obtained using a semantic map to predict $\mathbf{z}$ from fMRI signals, instead of original images as we did in the main paper. In this analysis, we used masks provided by MS COCO for semantic segmentation. These masks provide a semantic label for each pixel in each image, and it is thus considered that the original images are coarsely approximated. Interestingly, using a semantic map provides the same level of accuracy as using the original images despite there being much less information contained. Specifically, we found a slight increase in semantic fidelity when we used semantic maps instead of original images, although the visual similarity slightly decreased. The present results are exploratory and we leave a more detailed analysis for future work.

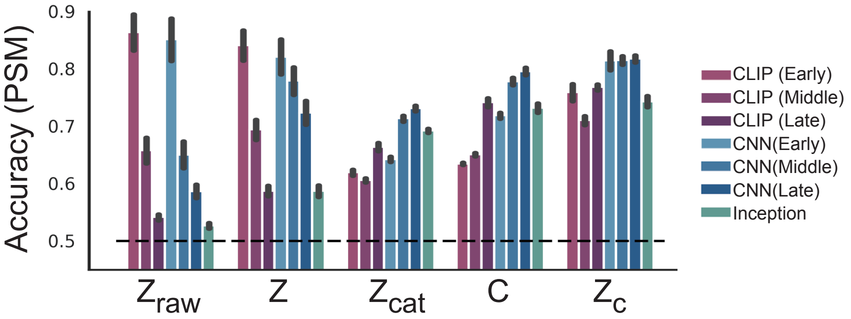


Figure B.2. Additional identification results

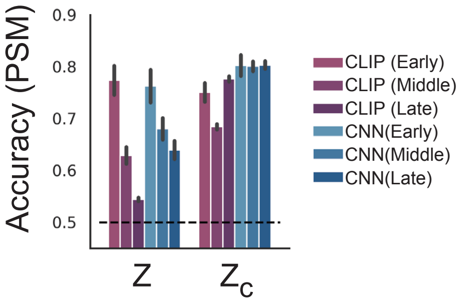


Figure B.3. Identification results obtained using a semantic map

#### B.3. Results for a Single Subject

We provide present additional results from for a single subject (subj01, Figure 3) in Figure B.4.

#### B.4. Results of All Subjects

We present additional results for all subjects (Figure 4) in Figure B.5.

### C. Additional results of encoding models

#### C.1. Results of a Single Subject

We present additional results of a single subject (subj01, Figure 6) in Figure C.6 for prediction accuracy with different noise levels to generate $\mathbf{z_c}$ from $\mathbf{z}$. We set the strength parameter for noising/denoising to 0.6 (low noise, top), 0.7, 0.8, and 0.9 (high noise, bottom). As expected, $\mathbf{z_c}$ with a reduced noise level injected into $\mathbf{z}$ produces a prediction map more similar to the prediction map obtained from $\mathbf{z}$.

#### C.2. Results of All Subjects

We present additional results for all subjects in Figures C.7, C.8, C.9, and C.10 for Figures 6, 7, 8 and 9. These results show that our results are robust across subjects.

#### C.3. Results from Different Random Seed

We present additional results for a different random seed in Figure C.11. These results show that our results are robust in terms of different random seeds.

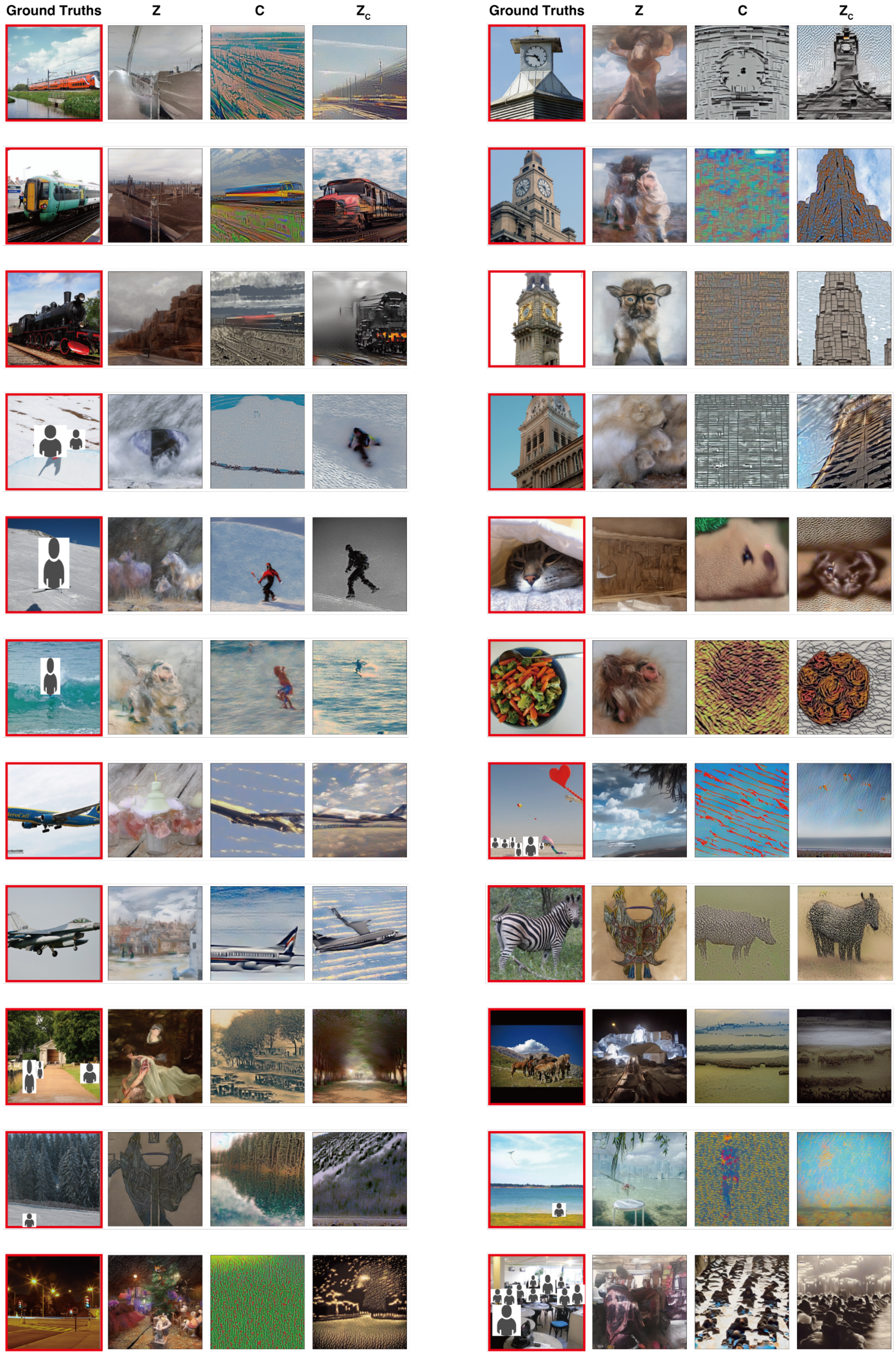


Figure B.4. Single subject results

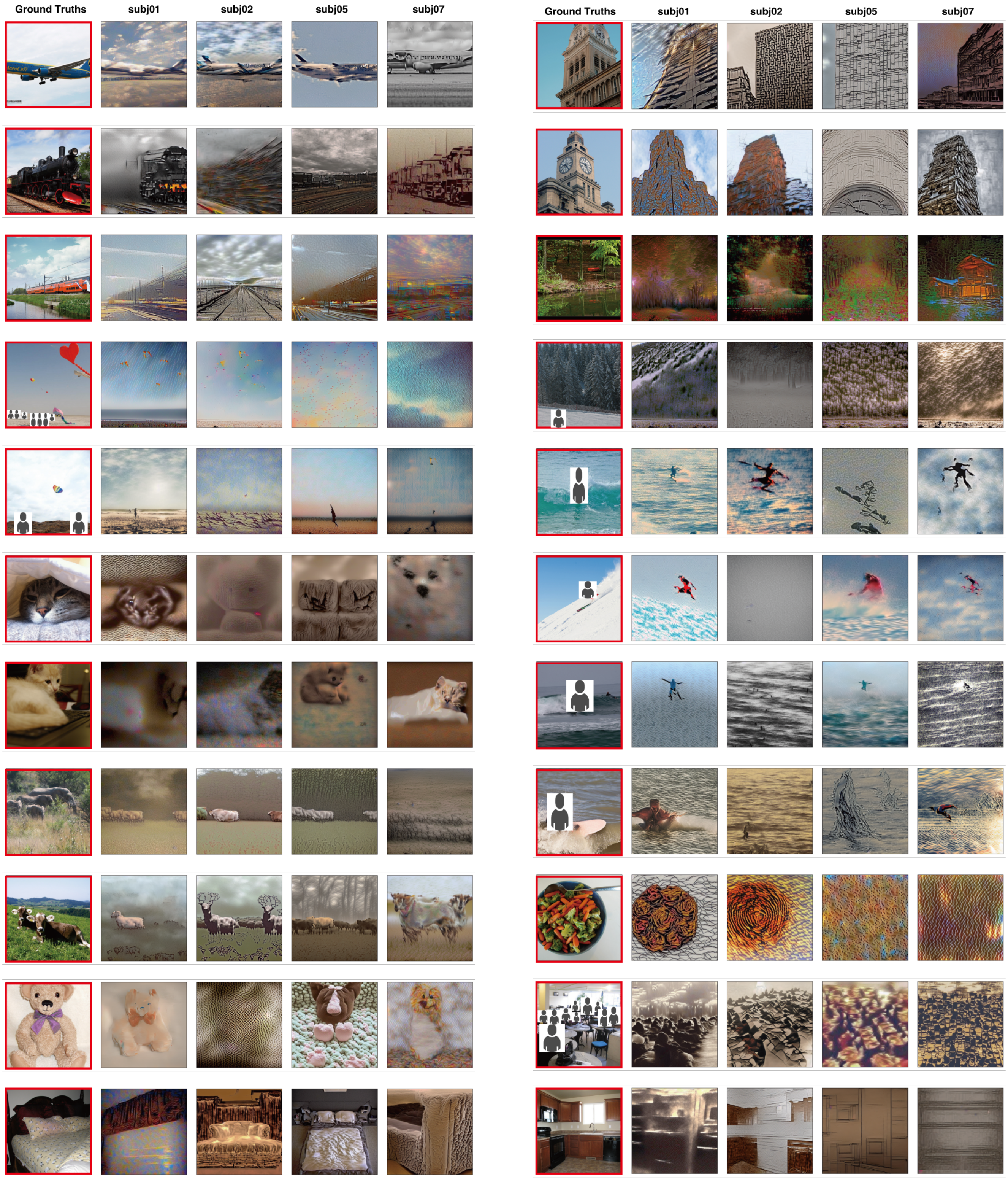


Figure B.5. All subject results

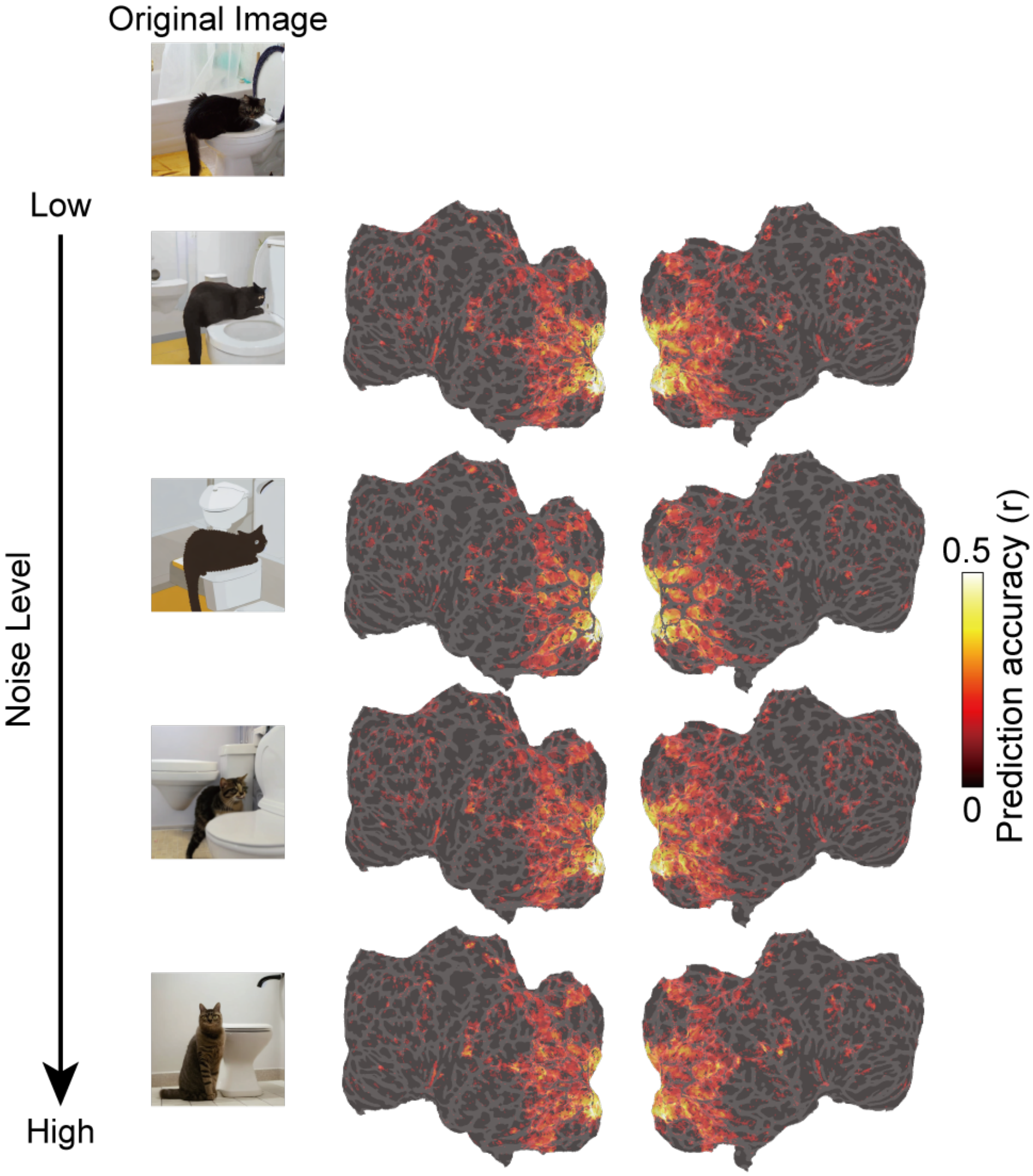


Figure C.6. Accuracy map of other noise levels for one subject

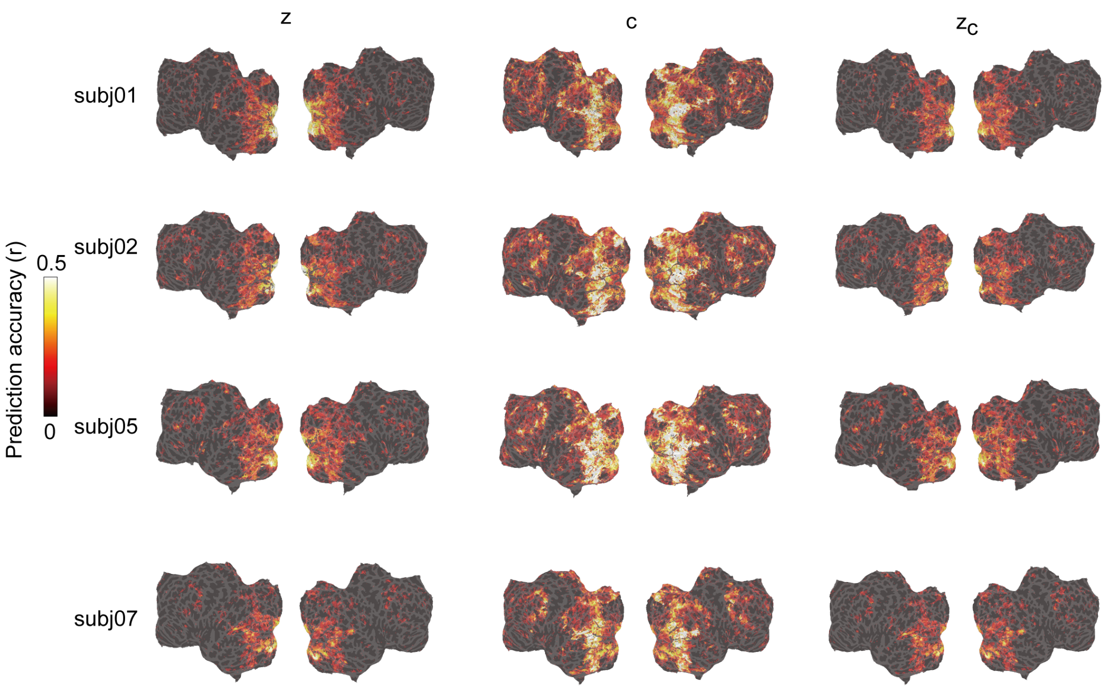


Figure C.7. All subject results for Figure 6

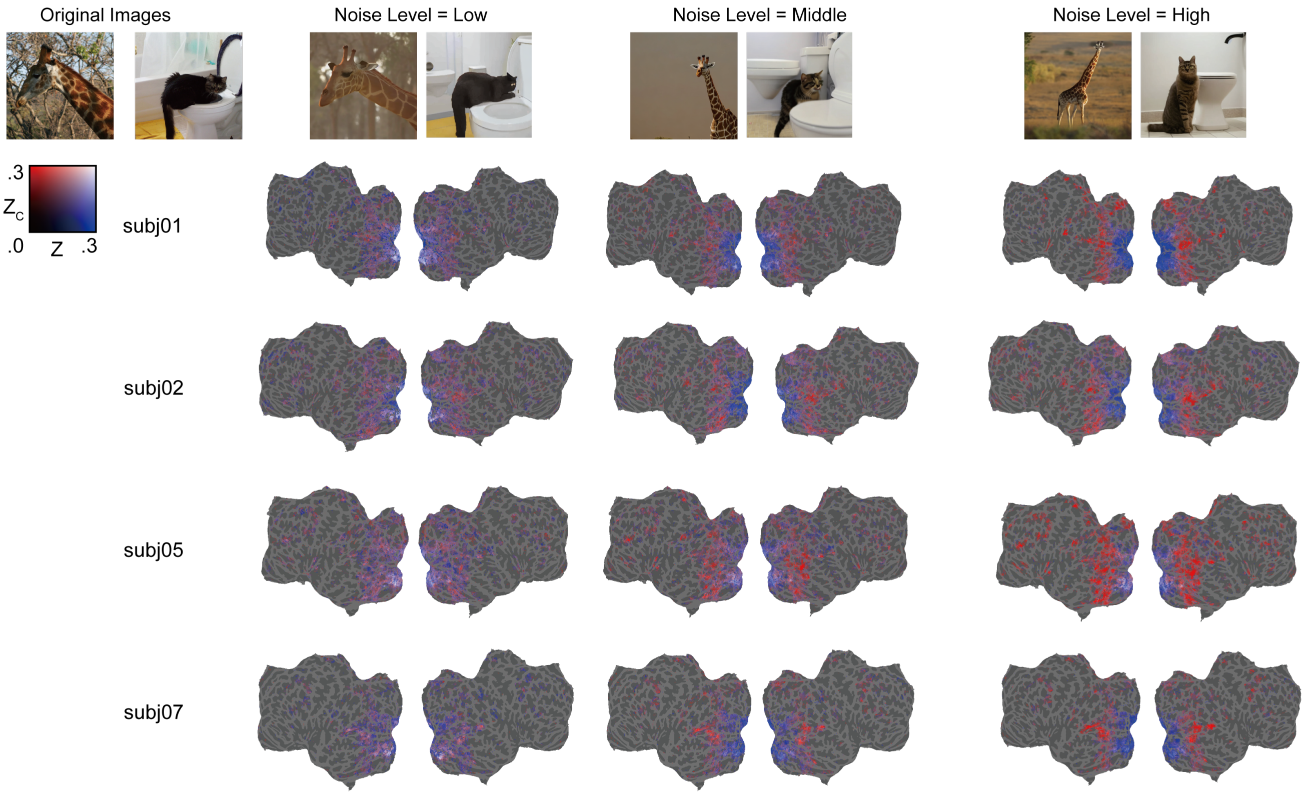


Figure C.8. All subject results for Figure 7

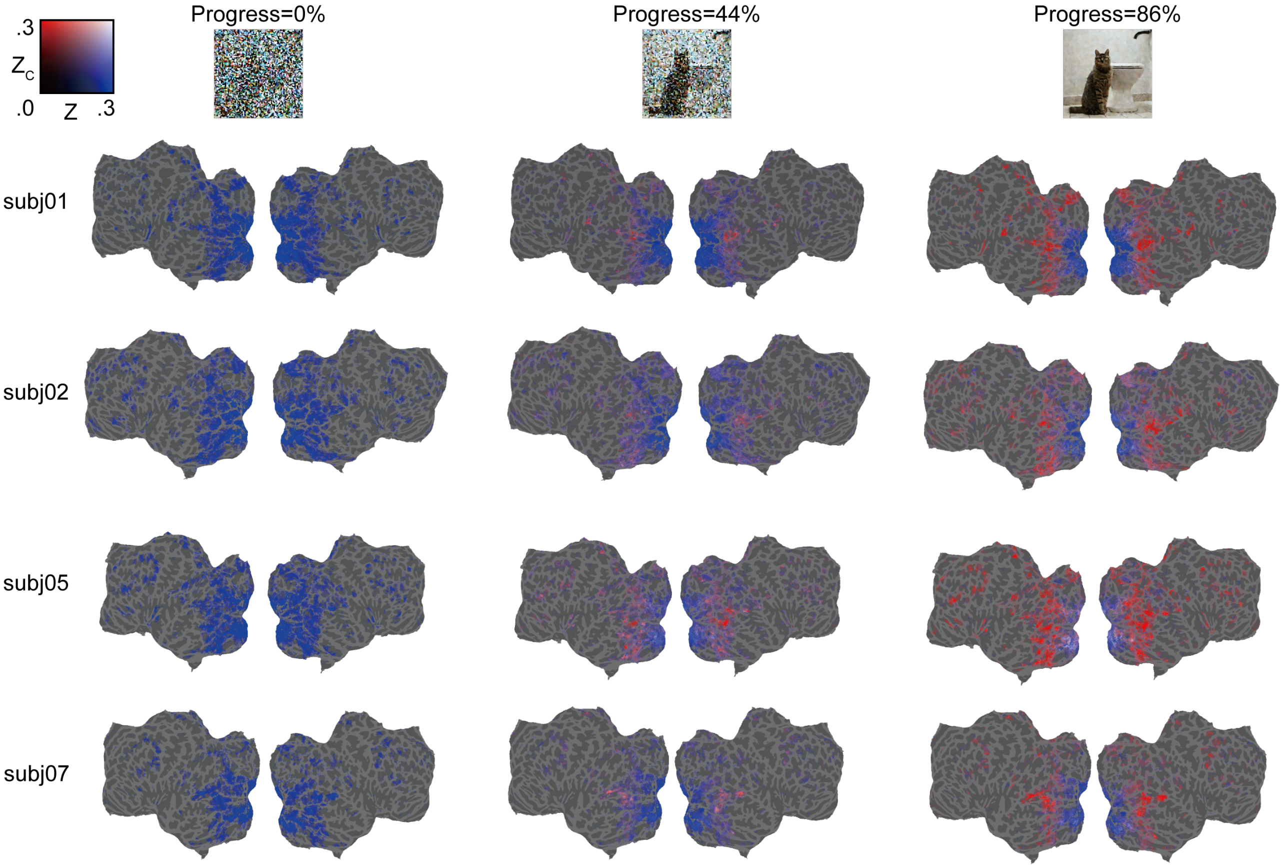


Figure C.9. All subject results for Figure 8

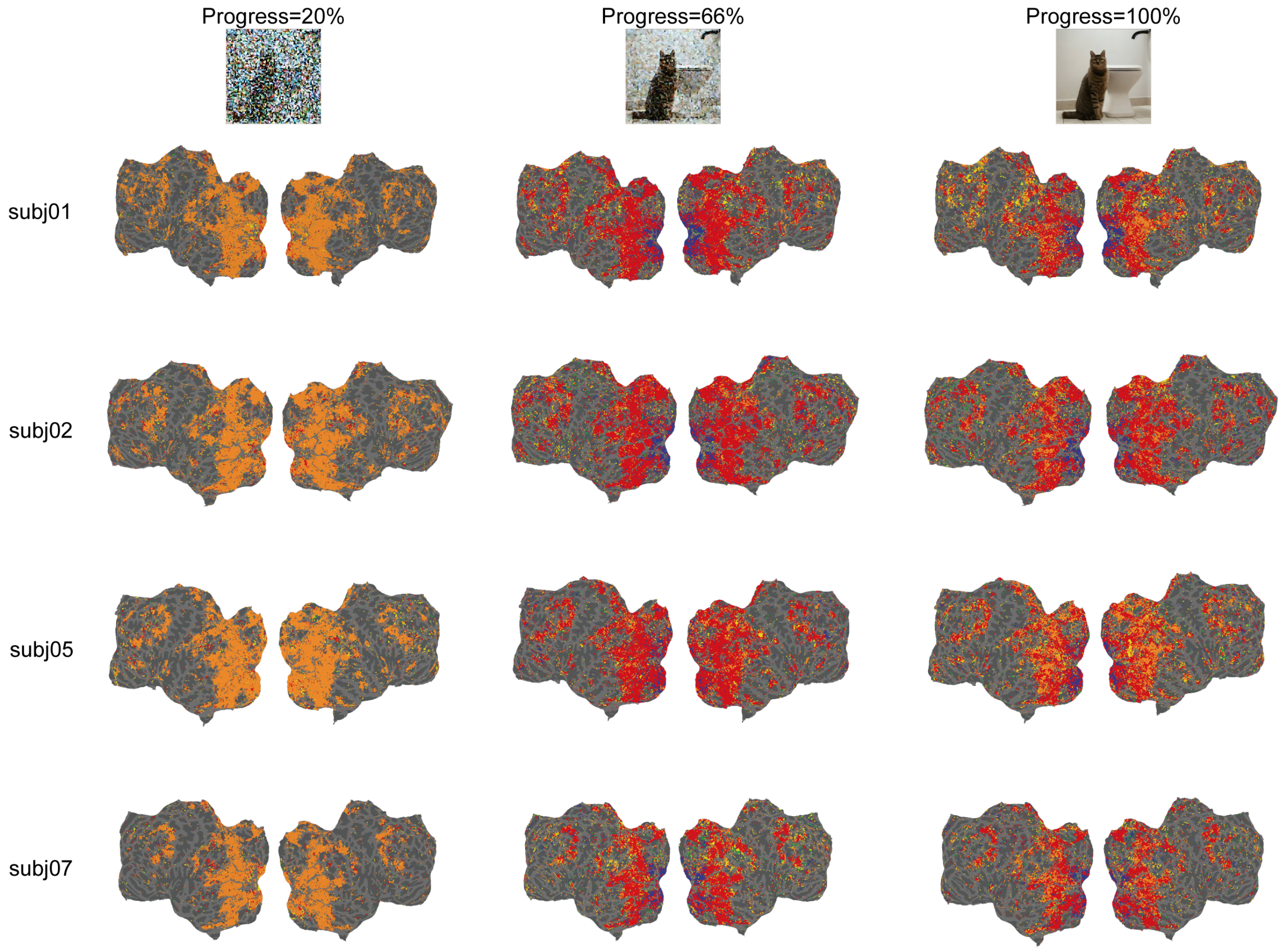


Figure C.10. All subject results for Figure 9

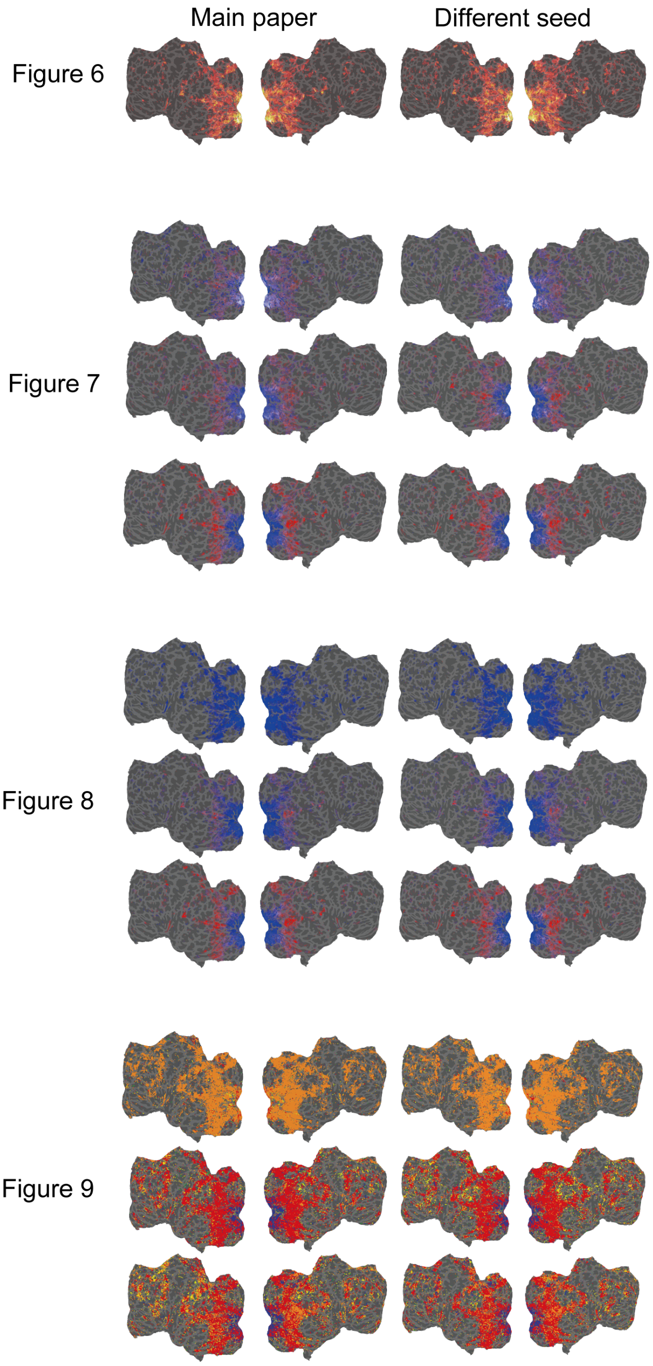

Figure C.11. Results for different random seeds
